# Spinning Disk Multifocal Microscopy for Dynamic Arbitrarily Shaped Surface Imaging at Centimetre Scale and Micrometre Resolution

**DOI:** 10.1101/2022.02.22.481397

**Authors:** Hao Xie, Xiaofei Han, Guihua Xiao, Hanyun Xu, Yuanlong Zhang, Guoxun Zhang, Qingwei Li, Jin He, Dan Zhu, Xinguang Yu, Qionghai Dai

## Abstract

The large-scale fluorescence microscopy has enabled the observation of dynamic physiological activities at the single cellular level across the mouse cortex, such as distributed neuronal population representations. However, video-rate high-resolution microscopy at sophisticated biological surfaces in nature keeps a challenging task for the tradeoff between the speed, resolution, and field of view. Here we propose Spinning Disk Multifocal Microscopy (SDiM) for arbitrarily shaped surfaces, which enables imaging at centimeter field-of-view, micrometer resolution and up to 30 frames per second across the depth range of 450 µm. We apply this technique in various microscopic systems, including customized macroscopic systems and the Real-time Ultra-large-Scale imaging at High resolution macroscopy (RUSH), in both the reflective mode and the fluorescence mode, and in the study of cortex-wide single-neuron imaging and immune cell tracking. SDiM provides an opportunity for studying the cortex-wide multi-scale cellular interactions in biological tissues.

## Introduction

Observation of cellular activities in multiple cortical regions is crucial to understanding a variety of biological processes, such as neural computation and immunological processes, but simultaneous imaging of the whole cortex at micro resolution is challenging. Recently, advances in skull window techniques have provided optical accesses to large areas of the brain. For example, the open-skull glass window techniques builds a curved glass or polymer optical window with the shape of a skull^1,2^, which enable large field of view (FOV) imaging across the whole cortex. The thinned-skull cranial window technique thinned the skull to reduce optical scattering^3^. Skull optical clearing techniques apply bio-compatible reagents to the skull, in order to remove the scattering particles and match the refractive index of the skull^4,5^. All above techniques have provided us unprecedented opportunities to investigate on the interaction between brain regions at the cellular resolution. On the other hand, high-throughput cellular imaging techniques has made cortex-wide recording possible. The Real-time Ultra-large-Scale imaging at High resolution microscope (RUSH) breaks the cameras’ barrier of data throughput, by employing 35 cameras simultaneously to capture centi-meter scale dynamics at the micro resolution and video-rate^6^. The two-photon light beads microscopy breaks the limitation of laser repetition-rate by breaking one strong pulse into 30 sub-pulses^7^. However, above-mentioned techniques are limited to a local brain region, or need to flatten the optical window. It remains impossible to image the cortical structures at cellular resolution due to the trade-off between the depth of field (DOF) and resolution of these methods. As the curvature increases at the border, a cortex-wide flat craniotomy window squeezes the brain tissue and alters the pressure on brain, which is a potential cause of hernia ^8^. So a microscopy with high resolution, large FOV, and high capture rate is required.

Several techniques have been proposed to expand the DOF. Reducing the numerical aperture (NA) increases the depth of field, but leads to low resolution and low signal to noise ratio (SNR), inducing photo-bleaching and phototoxify; Fast focal distance adjustment elements, such as electrically tunable lens and TAG lens, encounter the tradeoff among FOV, resolution and speed, which are thus inapplicable to high throughput lens^9–12^; Piezo stages under resonance frequency cannot achieve fast scan and high load simultaneously, which is also challenging to achieve accurate synchronization^13^; focal length expending techniques, such as SPED microscopy and cubic phase plate-base techniques, acquire larger field of view at the cost of lower SNR^14–16^; 3D imaging techniques, such as light field microscopy^17–21^, sacrifice the resolution for improved speed, but bring high computation cost. Besides, all of these approaches improve imaging speed but suffer from optical artifacts (especially beyond the native focal plane or point). Also, they are restricted to small sample depths in general, and require complex instrumentation and alignment procedures.

To this end, we propose the Spinning Disk Multifocal (SDiM) microscopy, which enables arbitrary-shaped surface imaging while maintaining spatial resolution at large FOV. At the core of SDiM microscopy is fast modulation of the focal plane, building upon the optical mechanisms that refractive index mismatch induce focal shift. Focal plane of the objective is altered by a fast-spinning disk with different glass thickness, rather than the moving specimen or objective. Together with the imaging area selection acquired by illumination area selection (using a digital micromirror device, DMD), this technique provides the capability to scan thirty frames in a second, which is limited by the camera. We demonstrate the capability of SDiM microscopy by imaging fluorescent microsphere samples at micrometer resolution, and by recording neural activities across the entire superficial cortex of transgenic mouse expressing GCaMP in microglia cells and neurons. This open-source cost-effective device can be easily adapted to a majority of microscopic systems, and the focusing algorithm interface is compatible with various algorithms including deep neural networks.

## Results

### SDiM operating principle

The Spinning-Disk Multifocal microscopy is developed from the combination of fast axial scanning and sequential illumination. In wide-field fluorescence microscope, the captured image from the camera is the summation of intensity of each time interval in its exposure time, while the intensity of each time interval is the product of illumination intensity and the fluorophore density. In our system, when the lateral position is scanned rapidly and sequentially, we keep illumination of in-focus areas on, and that of defocus areas off, so as to scan the entire surface in one exposure period, as shown in Fig. 1a.

**Figure 1.**
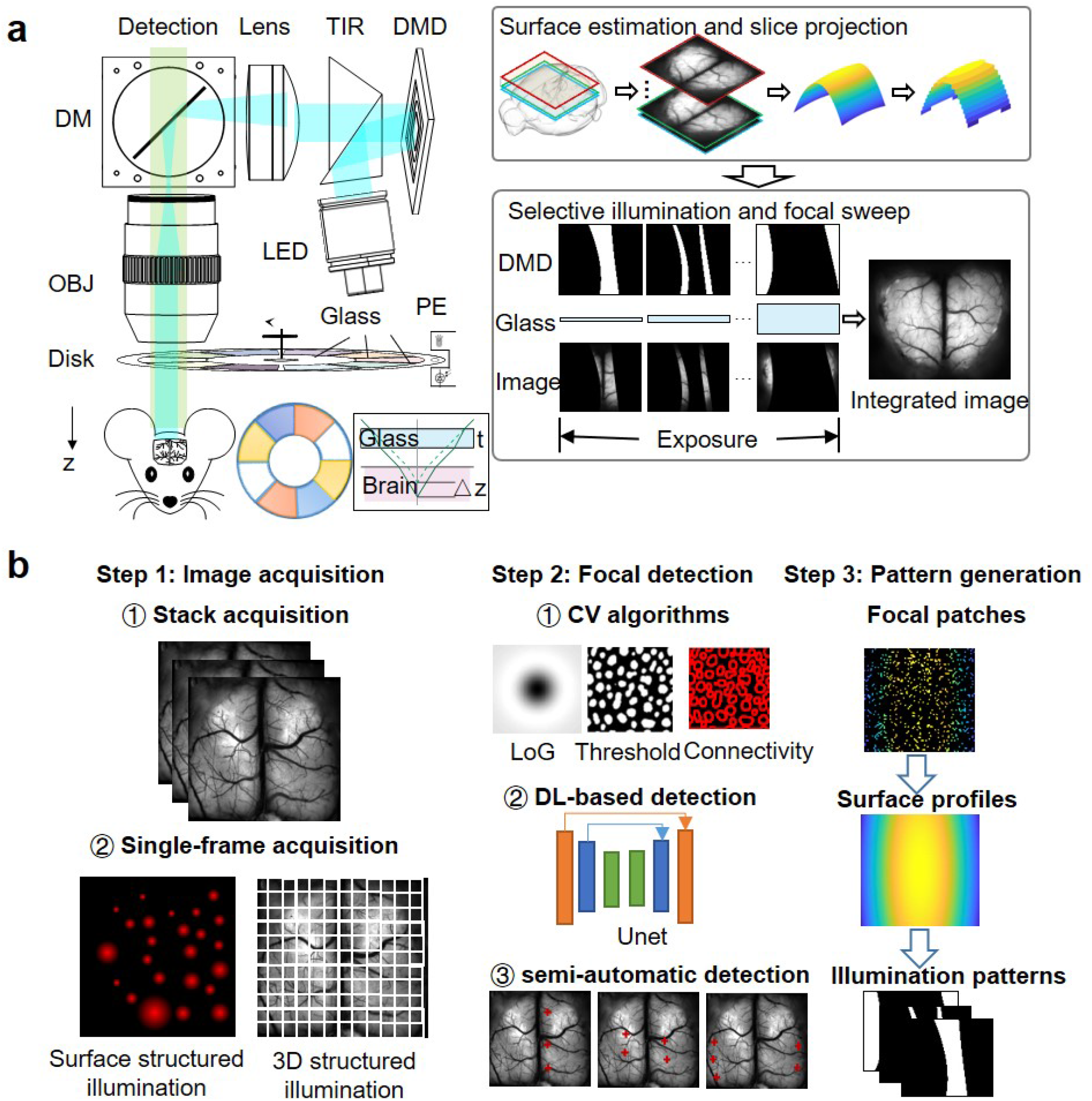
Schematic diagram of the system. a. Centre, the SPIM system is const of two parts, selective illumination and focal modulation. The selective illumination part consists of a digital micro-mirror device (DMD) conjugated to the image plane, which modulates the illumination patterns from the light source. The focal modulation part is a spinning disk with cover glass of different thicknesses, which shift the focal image to different depth. Left, Before the acquisition frame, a depth map is generated from previous measurement and prior knowledge, which is turned into an attention map of each layer and determines the DMD pattern of each depth. Right, in each exposure period, the spinning disk is rotate for one cycle, while at each depth position the DMD displays one attention map. The integrate acquired image is the integration of focused image from different depth. b. Several techniques to acquitted the attention map. The image acquisition could be a stack for static or slow varying surfaces or single frame for fast changing surface. Depth of each axial position can be determined by computer vision algorithms, and/or further refined by manual/semi-automatic detection algorithms. Deep learning algorithms can also be applied once we have enough dataset. Finally, we generate the depth map from the point cloud., and calculate the attention map of each layer.

The key element of implementing this approach is to extend the depth of field of the objective, while keeping the resolution and speed to image a large intact curved surface. While several related methods exist, including the use of an electric tunable lens and remote focusing, they alter the axial position of the objective by changing its focal length, but suffer from the tradeoff between the field of view, resolution and imaging rate. Here we deploy a high-speed spinning disk with coverglasses of various thicknesses between the objective and sample, thereby introducing different focal shifts at different thicknesses, as previously described in Ref.^22^ and also shown in Supplementary Figure S3. Building upon the fact that the refractive index mismatch in optical systems results in focal shift, this approach extends the depth-of-field by an order of magnitude while maintaining lateral extent of the point spread function. The position of the spinning disk is detected by an IR rotation encoder, which synchronizes the focal position and illumination pattern. When the focal depth is altered by the spinning disk, the digital micro-mirror device generates an illumination pattern sequence, in which each frame turns on the in-focus regions, and turns off the defocus regions, as shown in Supplementary Figure S4.

The illumination pattern is generated from surface profile, as shown in Fig. 1b. For slow-varying specimens, such as the brain tissues of head-fixed mice, we take an image or an image stack at different focal positions, and search for the focal regions at each depth. For continuously changing surfaces, we adopt a single-exposure focal algorithm: the entire surface is randomly and evenly divided into non-overlapping small patches, which are illuminated and detected at different depths, so that the surface profile is estimated from the in-focus patches, as shown in Supplementary Figure S11 and S12. We provide interference for different focal detection algorithms, for example, computer-vision descriptor-based algorithms (Laplacian of Gaussian^23^, morphology, *etc*.), deep neural network algorithms (U-net)^24–26^, and other manual/semi-automatic detection algorithms^27^. After the detection of focal patches, we stitch the surface profile with smooth function fitting, and discretize this profile into illumination patterns, as shown in Fig. 1b.

### SDiM microscopy enables micrometre-resolution and centimetre-scale imaging

To demonstrate SDiM’s advantages over conventional microscopes, we first compare their point spread functions (PSFs) of fluorescent microspheres on a spherical surface, generated from a simulated dataset. Conventional microscopes have to trade off between the lateral resolution, DOF, and light collection efficiency, as shown in Fig. 1a. Objectives with 0.3 NA achieve ∼ 1 µm resolution, but their depth of field is restricted; the peak intensity drops dramatically 10 µm away from the focus. Conversely, Objectives with 0.1 NA have 100 µm depth of field, but their full width at half maximum (FWHM) is 3 times larger than the 0.3 NA objective. Further, their peak intensities drop to nearly 1% of the high NA objective, so they requires large pixel size or high-excitation power to compensate for the photon loss, at the cost of reduced optical resolution and increased photo-bleaching.

We also simulate the same neural image taken from high NA and low NA systems at different focal depth, as shown in Fig. 1a. Compared with conventional microscopies, the SDiM simultaneously enables high resolution, large DOF and large FOV imaging by temporal multiplex: High resolution and high photon-efficiency is remained at different axial depth.

Then, we experimentally validated our predictions under the RUSH microscope with a 10×*/*0.3 NA objective. We tested our system’s parameters with 0.5 µm microspheres under cover-glasses of 10 different thicknesses, ranging from 0 to 1.6 mm. We plot the focal shifts of different cover glasses in Fig. 2b, which is linearly related to the glass thickness. We infer the refractive index from the focal plane offset to be 1.55, which agrees quite well with the ground truth (*n* = 1.52 for wavelength at 515 nm). Also, we show the lateral and axial PSFs are independent to lateral and axial positions in our field of view, which is one of our most significant advantages over conventional lens-based techniques. We plot the averaged FWHMs of microspheres, which exhibit uniform distribution except for the very corner. We also plot the distributions of FWHMs at different coverglass thicknesses, which indicates the PSFs do not increase significantly with the larger glass thickness.

**Figure 2.**
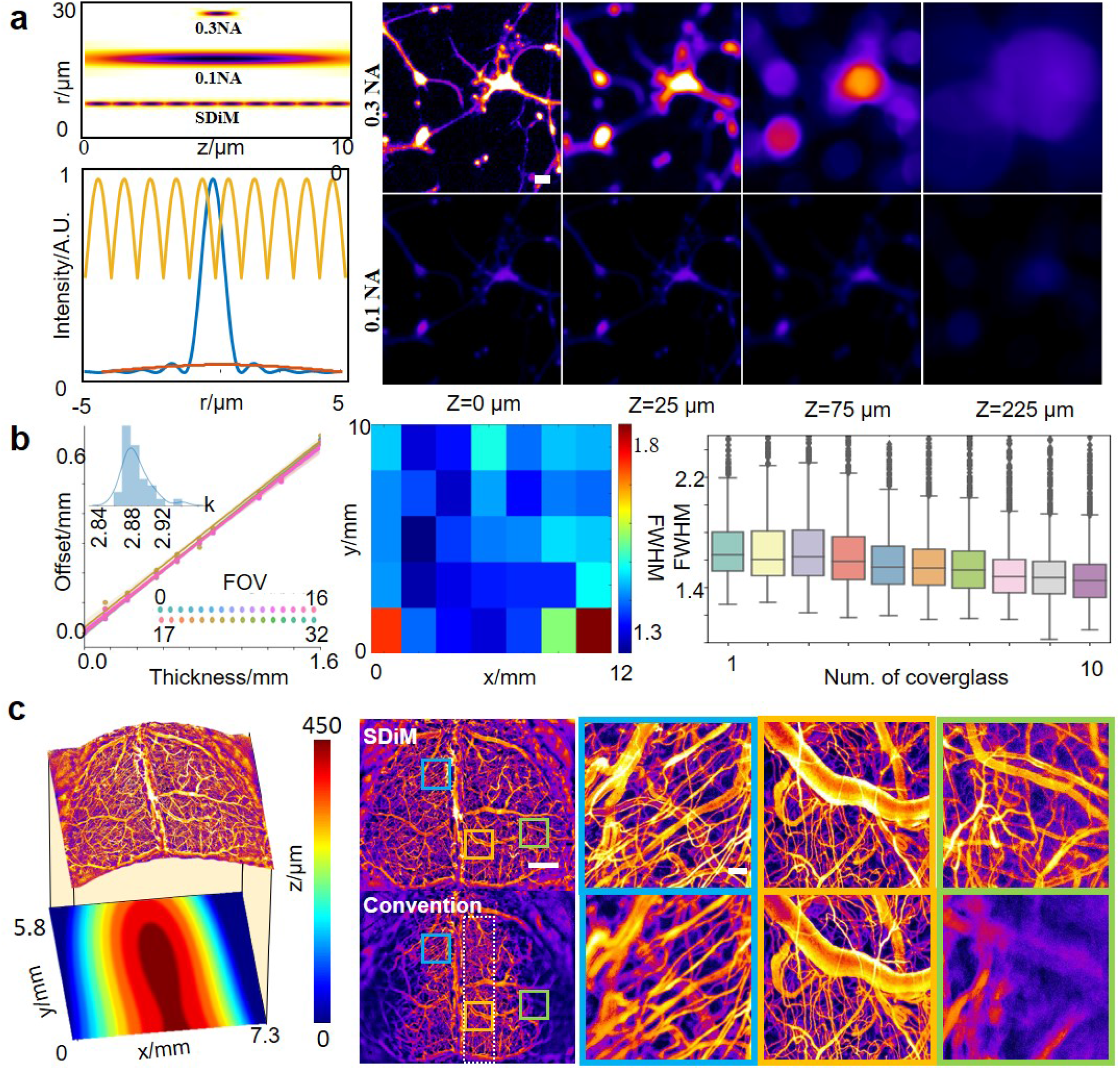
Parameters of the system. a. Theoretical simulations. Left top, point spread functions of different systems. High numerical aperture (NA) systems (0.3 NA) have high lateral resolution, but extremely small field-of-view (10 µm). Low NA systems (0.1 NA) have low lateral resolution, and modulate field-of-view (90 µm). Our SPIDM system have simultaneous high NA, large FOV. Their axial distribution is plotted in the left bottom figure. Right, simulation of a neuron with different defocus blur from 0 to 200 µm. For high NA the image quality is acceptable in a DOF of 30 µm, while for low NA systems the acquisition has significantly low photon efficiency for detection. As a result, we choose 50 µm as our depth step. 20 µm. b. Experimental validations on the Real-time Ultra-large Scale imaging at High resolution system. Center, Full Width at the Half Maximum (FWHM) is consistent across the 35 cameras in the whole field of view (except 3 in the very corner). Left, the focal plane of 32 cameras are shifted by the insertion of glass, with a slope of 3.8. This corresponds to a refractive index of 1.55, which matches the theoretical prediction of 1.52. Right, the later resolution of point spread function does not change with the insertion of glass with different thickness, from 0 to 16 mm. Scale bar, c. The brain image of a wide-type mouse with FITC injection in vessels. Left, the estimated brain image (top) and its depth map. Middle, later projections of the mouse brain images from SDiM (top) and conventional microscope with 0.3 NA. Three local zoom-ins are shown in the right, which shows conventional microscopes suffer from out-of-fucus blur, while SDiM enables globally all-in-focus imaging. Scale bar, 1 mm for the global views, and 100 µm for the local views.

Next, we assess the imaging quality in live mouse brains, so as to demonstrate the main advantages of fast and complete imaging of the intact superficial mouse cortex. We acquired image stacks from live, unparalyzed mouse injected with Fluorescein isothiocyanate (FITC)–dextran under a customized macroscope constructed from two Digital Single Lens Reflex lens. As demonstrated in Fig 2c and Supplementary video 1, the SDiM microscope allows cellular resolution imaging of the intact superficial brain, while conventional macroscopes focus on only the central region of the whole brain.

### SDiM microsopy enables rapid cellular-resolution functional imaging of the mouse brain

To demonstrate SDiM’s high-speed, cellular-resolution and large field-of-view imaging capabilities on complex surfaces, we sought to determine if it would be possible to capture fast micrometer-resolution spontaneous activity over the entire mouse superficial brain. We used the customized microscope to perform brain imaging of the awake Ai148D: rasgrf −2A-dCre mouse, as shown in Supplementary Video 2. This transgenic mouse strain expresses calcium indicator GCaMPf mostly distributed in Layer 2/3 in the superficial layer, which located 150 micrometer below the dura, and its cre recombination is regulated by trimethoprim injection. For a head-fixed mouse, its neural position is stable, so we take a time serial at each depth, and detect the three-dimensional neural position with morphology methods. Then we fit the surface with a second or third order polynomial function, as shown in the insets in Fig 3a.

**Figure 3.**
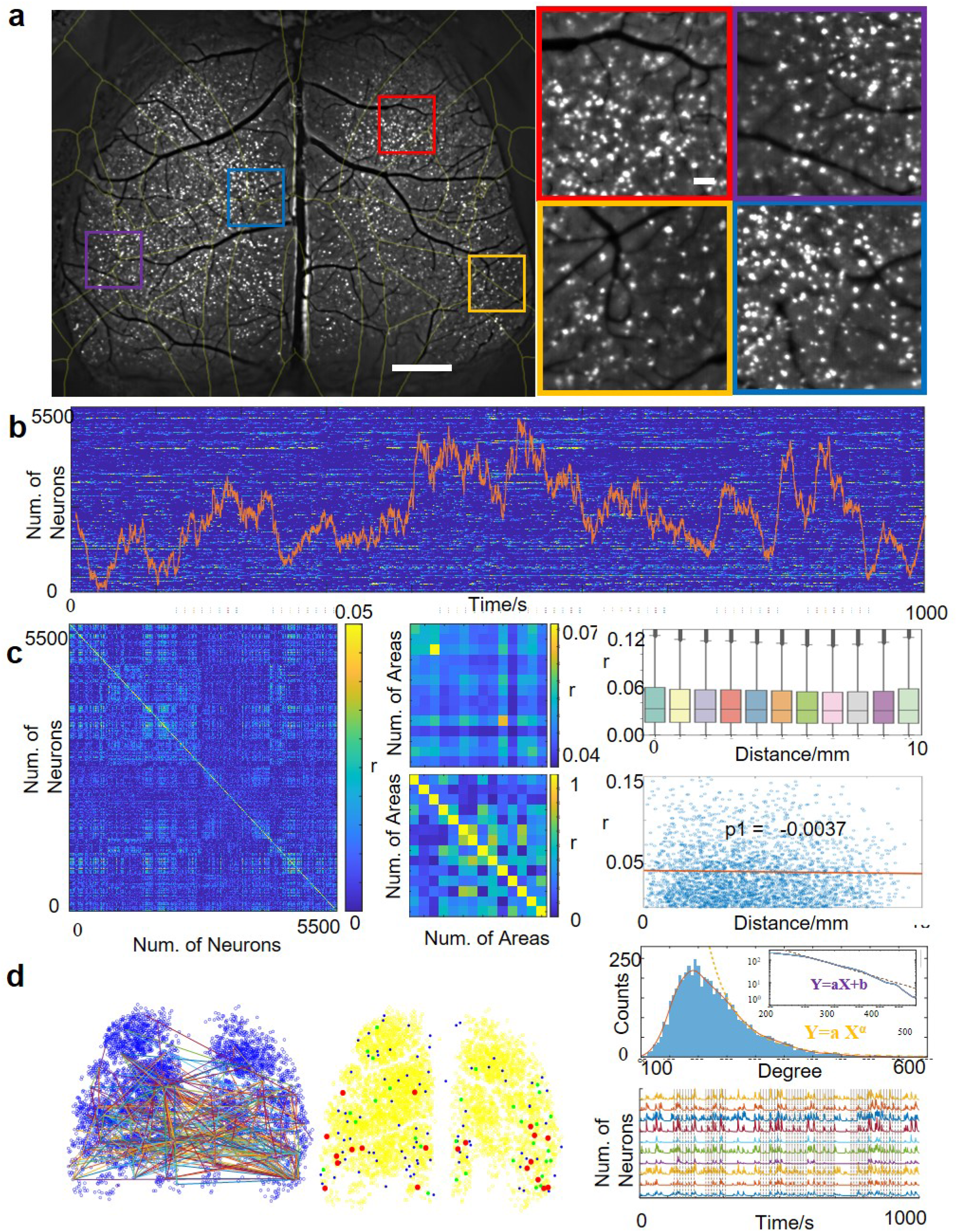
Images of a transgenic mouse expressing GCaMP6f in neurons. a. Top left, standard derivation of neural activity from our SDiM microscopy, with four zoom-ins across the FOV are displayed in the bottom. The image is registered to the Allen Mouse Brain Common Coordinate Framework. Top right, the constrained nonnegative matrix factorization (CNMF) algorithm detect 5400 neurons in SPiM, while only 1000 in the conventional microscope. Scale bar, 1 mm. b. Traces of the 5400 neurons, with their principle component and visual stimulus. c. Left, correlation of 5400 neurons sorted by brain regions, averaged to avoid aliasing. Middle top, averaged correlation of each brain region. Middle bottom, correlation of averaged neural traces of each brain region. Right, relationship between correlation and distance. Scale bar, 100 µm. d. Left top, edges in the graph, resulted from threshold correlations. Left bottom, Hub neurons in the network. Middle top, the degree of each neuron is fitted with the power distribution. Right top, traces of ten neurons with highest correlations. Right bottom, the hub neurons keeps relatively stable in time.

The entire superficial cortex is imaged in a snapshot, and SDiM detects 5,400 neurons via the constrained nonnegative matrix factorization algorithm. The imaging speed of SDiM is limited by the speed of current sCMOS cameras at up to 50 frames per second, but we set the speed to 10 Hz in order to match the dynamics of the calcium indicator. These neurons are distributed evenly across the whole field of view, as shown in Fig. 3b. In contrast, the conventional microscopy with 0.3 NA detects only 1,000 neurons in its depth of filed. We further zoomed in four selected regions in Fig. 3a, where the contours of all somata are visualized at high resolution. In contrast to the COSMOS microscope^28^, SDiM achieves single-cellular resolution, benefiting from the high optical resolution and high light efficiency resulting from 0.3 NA and 3 µm sampling rate.

The high throughput of neural detection enables large scale network analysis from single neurons. We plot the neural traces and their first principle component in Fig 3b, as well as correlations of the 5400 neurons (Figure 3c left, average filtered to avoid display aliasing). The cellular correlations show much more complex structures, which is quite different from the brain-area averaged correlation and the correlation of averaged brain region activity shown in Fig. 3c middle. To test whether the distance between neurons is the dominant factor in their correlations, we plot all neural correlations versus neural distance in Fig. 3c right. Unlike brain-area averaged activities, it is shown that the neural distance has little impact on single neural activities. We further study the network properties in neural dynamics. As demonstrated in Fig. 3d, we constructed a graph by thresholding neural correlations, and plotted the neurons with the highest degree in Figure 3d left. We defined the degree as the sum of absolute value of neural correlations, and plotted the statistics of nodes with different degrees. As shown in Figure 3d, the degree of nodes obeys the power law, as described in the small-world network, and the hub-neurons have similar patterns over time. We also demonstrate the wide scope of applications by imaging neural activities in a mouse injected with adeno-associated virus expressing GCaMP6s (Supplementary Figure S6), and in a cleared skull (Supplementary Figure S5), in which the neural activities are visualized across the field of view.

### SDiM microsopy enables real-time tracking in vasculatures and in the superficial mouse brain

To further validate the application of SDiM in *in-vivo* cellular imaging, we perform immune cell tracking in a B6/C57 mouse implanted with a crystal skull. To labels neutrophils and a subgroup of other immune cells, we performed a intravenous injection of Ly-6G^29^, 2 hours after the craniotomy. Different from the neural imaging, these cells locate at the superficial of the cortex, so their axial position were located the assistance of superficial vessels.

We demonstrate SDiM has the capability of capturing events of different time scale in Supplementary video 3. Immune cells have different depth across the whole superficial brain, as well as distinct dynamics. In vasculatures, the immune cells flow with blood at the time scale of sub-second to seconds, while the neutrophils migrate on the surface at the seconds to tens of seconds scale, as shown in supplementary video 3. In Fig. 4b, we show a large cell travel through a vessel in seconds, and in Fig. 4c we find a cell migrate from one vessel and move towards another vessel in 2 seconds, and then flows away with the blood. We also track the single-cell activity with Trackmate^30^ and plot the long tracks of immune cell in Fig. 4d. The mean squared distance of these tracks is distinct from the random Brownian motion, which indicates there are potential communication or coupling among the cells.

**Figure 4.**
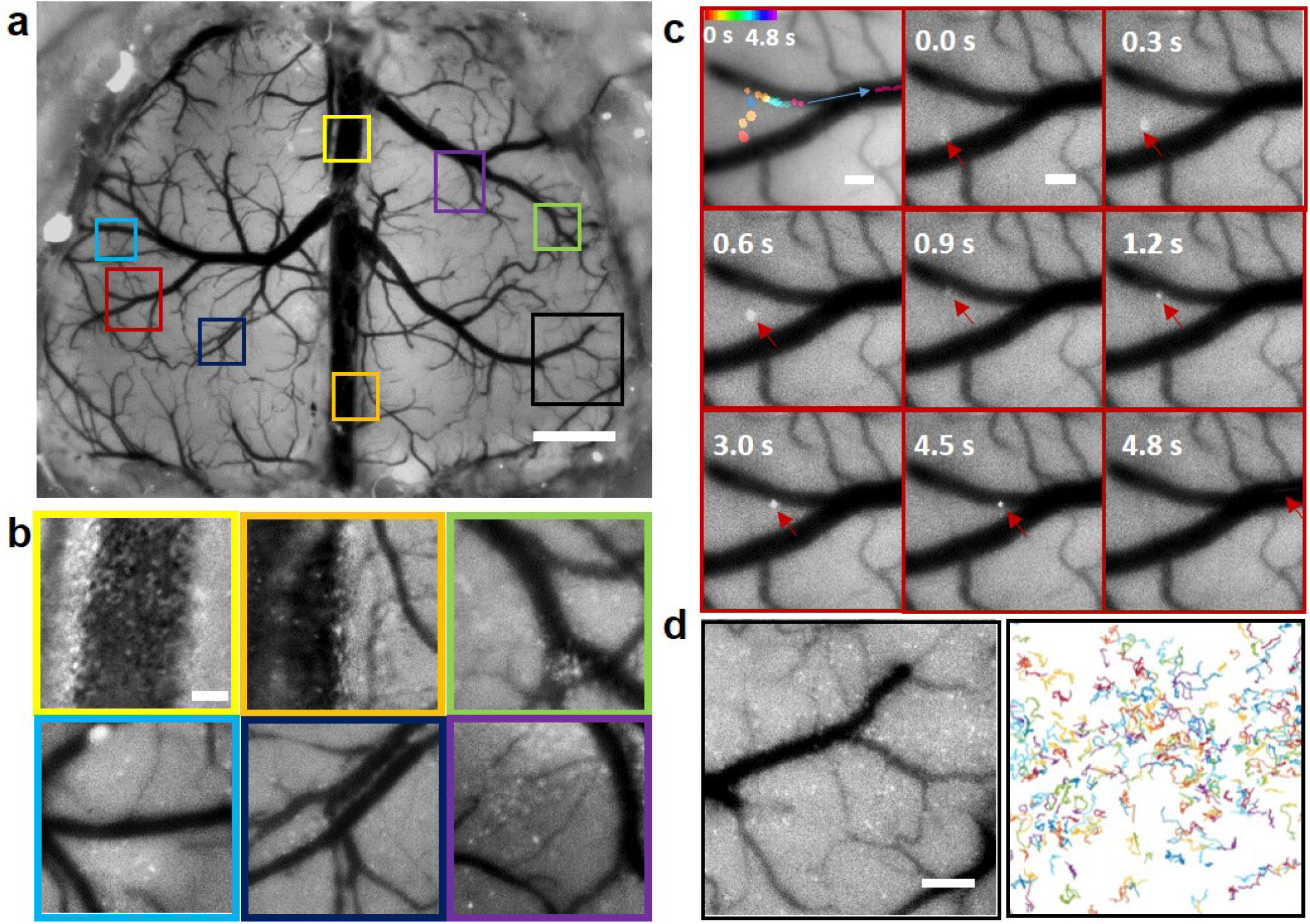
Recording of neutrophil transportation in a mouse brain after craniotomy. a. A snapshot of the neutrophil across the whole mouse brain. Scale bar, 1 mm. b. Zoom-ins of areas across the whole mouse brain in Fig. Our system is capable of simultaneous capture of fast dynamics in vessels and slow motions activities on the superficial brain. Scale bar, 100 µm. c. One zoom-in of areas in Fig. a (up), and the neutrophil traces are plot in the bottom.Scale bar, 100 µm. d. A time eclipse sequence of a zoom-in view in Fig. a, in which an immune cell migrates from one vessel to another vessel in seconds. Scale bar, 200 µm.

The capability of simultaneous capture slow events (in brain tissues) and fast events (in vessels) indicates our method has wide applications in immunology. To capture the fast events of single cells, we need high speed, high resolution and large field of view simultaneously. Further, immune cells are vulnerable to heat^31^, so it is important to reduce the photobleaching and phototoxify in imaging. Compared with conventional axial-scan microscopes, SDiM microscopy masks 90% of the defocus illumination, which is crucial for the normal behavior of immune cells. We also show the images of microglia cells in a CX3CR1-eGFP mouse in in Supplementary Figure S7 and S8.

### SDiM microscopy enables real-time microscopic imaging on complex surfaces and on the reflective mode

Finally, we demonstrate SDiM microscopy is able to image microscopic dynamics on other complex surfaces. We dropped FITC solution on the petiole of a fresh Epipremnum aureum leaf, and took a time elapse sequences of solution transportation in veins. First, we determine the surface profile of the leaf, as shown in Fig. 5a. The images on the complex surface is taken at 5 Hz. Compared with conventional focal-stack acquisition, only 10% images are taken with much less data storage. We plotted the results of both SDiM and the conventional microscope in Supplementary Figure S9 and in Supplementary video 4.

**Figure 5.**
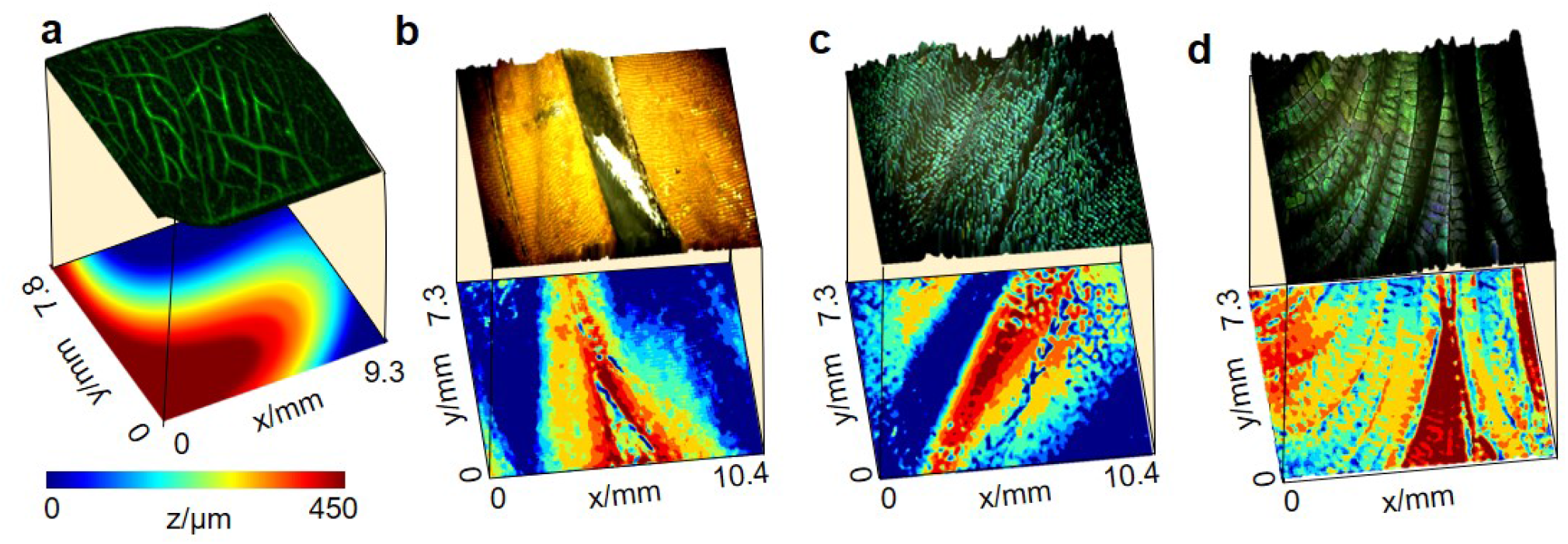
SDiM images of complex surfaces. a. Image of a scindapsus aureus leaf with FITC solution. Left, the transportation of FITC solution in veins is all-in-focus in SDiM, while blur in conventional microscopes. Right, a 3d projection visualize the peak time of each vein, while their depth map is shown below. b. Wings of butterflies with their depth map.

To further demonstrate the SDiM’s applications in different systems, we present images taken from a reflective microscope, with white light illumination and RGB camera acquisition. We took the images from the wings of three different kinds of insects. Due to the micro structures on the wings, conventional microscope can hardly image the entire field of view with high resolution, while our method is able to take images at different depths in a snapshot. This application indicates our method possesses high potential in many real-world applications, such as biomedical sample diagnosis.

## Discussion

Building upon the opportunities provided by the development of skull window techniques and optical cellular-activity sensors, we have developed the SDiM microscopy, which combines the extended depth of field with the large field-of-view microscopy to provide high imaging speed across a complex surface (such as the intact superficial mouse cortex) at cellular resolution. SDiM microscopy provides high spatial resolution comparable to standard epi-fluorescence microscopes, while providing the potential to scan more than 30 volumes per second, as a DMD can change its pattern in less than a millisecond. The speed of this method is therefore only limited by camera acquisition rates and rotation speed of the spinning disk. Compared with other volumetric imaging techniques, such as light field microscopy and the Spherical-aberration-assisted Extended Depth-of-field microscopy, our method has the highest light collection efficiency, and no requirement for deconvolution. This approach can also be extended to more specialized systems. In Supplementary Figure S10, we also present the design of SDiM in a high-resolution system with 1.45 NA, in which the spinning disk is inserted in the image space.

By choosing the appropriate combination of cover glass number, glass thickness, and detection objective NA, SDiM can achieve the desired depth of field; for example, we demonstrated micron resolution structural imaging in fluorophore beads at 1 mm depth. Three factors need to be considered to decide the glass thickness interval: the first is numerical aperture of the objective *NA*, pixel size *p*, and character feature size of the specimen *d*. For example, our system has 0.3 NA, and the pixel size is ∼ 0.8 µm and 3.2 µm respectively for RUSH and customized system, while the diameter of mouse neural soma is about 15 µm. As a result, the theoretical depth of field of an ideal point source is derived by *L*_1_ = 2*λ/NA*^2^ ≈ 10 µm. However, considering the shape of the features, the areas of the somata will double when the distance is *L*_2_ = *d/NA* ≈ 50 µm. So we choose the minimal glass thickness as *nL*_1_*/*(*n* − 1) ≈ 30 µm and the maximum glass thickness difference according to *nL/*(*n* − 1) ≈ 150 µm. Here we choose the thickness interval as 50 µm. Then the number of cover glasses is calculated by the focal interval *N* = 450*/*50 + 1 = 10. The excitation power needs to be increased by N accordingly to compensate for the reduced illumination time. Correspondingly, the SDiM illumination power is roughly N times of the wide field illumination, but most energy is reflected by the DMD and does not cause sample heat. Only the in-focus regions are illuminated, so the photobleaching and photodamage is reduced by a factor of N, compared with 3D scanning. The cost of increasing the illumination is not high: for example, it costs less than 1$ to get a 3W blue LED., this scaling effect can be relieved by illuminating several depths simultaneously close to the focal plane, at the cost of slightly reduced lateral resolution. Another alternative is to replace the amplitude-modulating DMD with a phase-modulating spatial light modulator, which can fully utilize the excitation power but at a higher cost.

To evaluate the effect of additional cover glasses to the optical system, we simulate the point spread function of system and enclosed energy with Zemax, with the glass thickness varying from 0 to 30 mm (corresponding to focal shift from 0 to 10 mm), under the condition of 0.3 NA. As previously described by the COSMOS microscopy, the full width at half maximum will remain unchanged with the increase of glass thickness, but the energy disperse significantly with the increase of radius. As shown in Supplementary Figure S2, if we require the capsuled energy in a 10 µm diameter circle is greater than half total energy, a maximum coverglass thickness of 15 mm is alowed for our SDiM system, which corresponds to 5 mm focal shift. Next, we consider the mechanic errors in manufacture, as shown in Supplementary Figure S1. If we allow the lateral shift to be the diameter of neural somata (15 µm), we can accept a tilt of 17^°^. This precision can be easily satisfied even with UV-glue. Further, if we consider the overlapping illumination regions, its width is about 20 µm, which is far smaller than the full field of view. As a conclusion, the additional coverglass in SDiM remains the high image quality of the optical systems.

Enabled by the capabilities of SDiM, we have observed cortex-wide neural dynamics in the awake mouse brain, and neutrophil dynamics after the craintomy. This indicates the SDiM method is particularly useful to neuroscience and immunology researches, for its low cast and easy implementation. Furthermore, we have demonstrated this method applies to other complex-shape surface such as leaves of an epipremnum aureum and wings of butterflies, thus could have a wide variety of applications including clinical diagnosis. Considering the wide applications of spinning disk, such as the color filter wheels in laser projector, this method also has the potential to be applied in the stereo display. In conclusion, together with advancing imaging processing algorithms, SDiM enables high-throughput, high speed, large-FOV, high-resolution imaging, thus paving the way to arbitrary surfaces dynamic recording in life science applications.

## Methods

### SDiM Implementation

One SDiM microscopy prototype was built on the previously described RUSH^6^ backbone, which provides an efficient platform for testing the point spread function and field of view with the insertion of the spinning disk. Figure 2 shows the details of SDiM optical implementation. Briefly, SDiM microscopy is generated from high-speed focal modulation and synchronized dynamic illumination area selection. High speed focal modulation is enabled by a customized aluminum spinning disk, as shown in Extended figure. 9 cover glasses with various thickness ranging from 0.17-1.5 mm is fixed to the spinning disk with UV-cued glue, and stabilized with tapes. A paper slit is attached to the edge of the disk to monitor the speed, with the help of a pair of IR emitter and detector. The spinning disk is fixed with a rotation stage, with a maximum rotation speed of 100 fps, and the rotation speed could be controlled by DC voltage. An optional cage could be added to secure the system. The spinning disk is placed between the objective and the specimen. Dynamic illumination region selection is enabled by digital modulation device conjugated to the image plane, with a maximum rate of 10M Hz. Each coverglass on the spinning disk corresponds to a specific region on the specimen, and is mapped into a DMD region. The excitation light source (LED, central wavelength 488 nm filtered by a bandpass filter) illuminate on the DMD, and the light beam passes scan lens, dichroic mirror, objective lens, and the spinning disk to illuminate the specimen. In the RUSH configuration, the emitted signal is detected with a camera-array-based wide-field detection arm, including a detection objective, a dichroic mirror, an emission filter, a tube lens array and an sCMOS camera array (Microvision, China). Details of these parts were described previously^6^. Another SDiM microscopy prototype was built on a customized microscope. We used low-cost second-hand Digital Single Lens Reflective (DSLR) cameras lens as the objective and the tube lens, and the image is acquired by an sCMOS camera. For non-fluorescence applications, the light source is replaced with a white LED light and the detection device is replaced with an RGB camera.

### Signal Synchronization

A DAQ device (National Instrument, USB-6363) is used for synchronization. For RUSH configuration, the sCMOS cameras are internally triggered, so we applied PID control to modulate the input voltage of disk, and used the frequency of spinning disk and camera output trigger as input. The camera output trigger is generated at the beginning of each frame, and the frequency of spinning disk is calculated from the signal from IR receiver. Then the IR signal is used to drive the DMD patterns. For the customized microscope configuration, position trigger of the spinning disk is taken as the input signal to drive both the start of camera exposure and DMD transition.

### DMD and imaging calibration

Single pixel images are projected on the DMD and acquired by the camera. Then the coordinates of the brightest pixels are read. Then we apply geotrans function to the coordinates of these five image pairs, which returns the transformation between camera pixels and DMD pixels.

### Surface Fitting

The first step of surface fitting is to detect the in-focus regions of the image. Here we offer three alternatives to detect the in-focus region: 1. computer-vision based methods based on gradient operators, such as the standard derivation on time and the Laplacian of Gaussian on spatial coordinates; 2. Human-invention methods that manually select the region of focal area; 3. Deep-learning methods. Different methods could be combined for different applications. Input image could be a stack of images illuminated at different height, or a single specially-designed pattern. For the blood vessel imaging, we applied a single mosaic image of different height illumination, and extract the regions with highest contrast. After the discrete regions are selected, a smooth or pre-defined surface shape could be applied to fit the surface. Then the height of the surface is discrete according to the glass thickness and this surface is divided into regions of different height. DMD masks of different heights are generated from these regions.

### Simulated PSF generation

The PSF of systems with different numerical apertures is generated with codes adapted from equations from Ref. Convoluted images of neurons is generated from the 2D convolution of a cultured image from the RUSH system and xy cross-sections with different system parameters. Here the assumption is that the axial resolution of the system is larger than the axial extends of neurons.

### System point spread function measurement

The point spread function of the system is measured under the RUSH system with a fluorescent microsphere specimen. The diameter of the beads is 0.5 µm, and about 10,000 beads are detected by our algorithm. We insert the spinning disk between the objective and our specimen, rotate the spinning disk to different cover glass positions, axially scan the stage, and simultaneously take the images from 35 cameras. Measurements of the point spread functions are taken from these PSF results.

### Animals

For this study, we used adult mice from the Jackson Laboratory. For blood vessel imaging experiment, C57BL/6J mice are injected with Fluorescein isothiocyanate-dextran (Sigma, MW 8000) before imaging. For microglia observation experiments, we used CX3CR1-GFP mice. For transgenic neural activity observation experiments, Ai148D:rasgrf-dCre mice are crossed from Ai148D and Rasgrf-dCre mice, and sparsely labelled adult C57BL/6 mice were made with 10 injection of a mixture of diluted AAV2-9-hSyn-cre and AAV2-9-Ef1a-DIO-GCaMP6f viruses (from BrainVTA Technology).

### Craniotomy

Chronic craniotomy was performed according to the procedures described in a previous report in accordance with the guidance of the Animal Care and Use Committees of Tsinghua University. We performed craniotomy with a window size of ∼8 mm ×8 mm, then installed curved optical windows and cemented the custom-made coverslips and aluminum head posts to the skulls.

### Imaging Experiments

For brain imaging experiments, the mice are head-fixed and awake during the imaging process. For visual simulation, a screen is placed before the eye of the mouse and grating of four different directions are generated from the matlab PsyLab toolbox. For sensory input, a small stick is attached to a galvanometer, and touch the whisker of the mice when stimulation is applied.

### Image Analysis

For the vasculature imaging, we take the standard derivations of the time sequence, and enhanced the image contrast with CLANE algorithm from imageJ^32^. For neural images, the neurons images are taken from the standard derivation of a time -filtered background-subtracted time sequence. The neuron positions are extracted from the constrained nonnegative constrained matrix factorization algorithm (CNMF-E). The neutrophil images are firstly filtered by a gaussian filter on time, and then subtracted a spatially filtered background of the averaged image. Trace of neutrophils are extracted with an ImageJ plug-in Trackmate^30^.

## Author contributions

H.X. conceived the idea of this project. X.H. designed and manufactured the system, with the assistance of H.X. X.H. developed the image processing framework and analyzed all the data, which the assistance from X.H., Y.Z. and G. Z.. G. X. and X.H. performed the biological experiments, with the assistance of HY.X.(craintomy), QW. L.(curved optical window), J.H. (neutrophil labeling), and D.Z.(skull-cleaning), under the supervision of XG.Y.. H.X. wrote the paper with editorial input from all authors. H.X and Q.D. supervised all aspects of the work.

## Acknowledgments

This project is supported by the National Science Foundation of China (62088102, 61831014, 61971256, 62171254). H.X. thanks Prof. Peng Xi for his suggestions to improve the manuscript.

## Additional information

The authors declare no competing interests.

